# A Mechanistic Account of Dedifferentiation in the Aging Visual System

**DOI:** 10.64898/2026.11.07.764836

**Authors:** Akilles Rechardt, Gabriele Bellucci, Robert M. Mok

## Abstract

To uncover the mechanisms of age-related dedifferentiation, we simulated neural degradation in a computational model of the ventral visual stream. Severing connections or adding noise, whether in inferotemporal cortex (IT) or upstream, reduced IT differentiation, indicating that dedifferentiation need not originate locally. Degrading peripheral V1, which preferentially drove place-selective IT neurons, selectively reduced place differentiation, a candidate mechanism for the stronger place dedifferentiation reported in aging.

## 1 Main

Age-related neural dedifferentiation is the reduction in the distinctiveness of category-specific activation patterns in older adults^1^ and has been linked to decreased memory performance^2^. Previous studies have shown reduced activation pattern differences between categories like faces and places in older adults’ higher-level visual areas^1–11^, including inferotemporal (IT) areas and identified candidate biological correlates of age-related dedifferentiation including neurochemical changes^12^, white matter degradation^13^, or gray matter volume loss^10^. However, despite well-established evidence for dedifferentiation, whether these forms of neural degradation are mechanistically *sufficient* to produce dedifferentiation in a network and how specific forms of neural degradation (e.g., neural noise, synaptic connection loss) cause it remain unknown. A second, more specific phenomenon also remains unexplained: dedifferentiation may not affect all categories equally. The parahippocampal place area generally exhibits more consistent age-related dedifferentiation for places or scenes than other category-selective areas do for objects and faces^3–9^ (although see^1,10,11^). Some have suggested gaze differences^3^ and lifetime experience^14^ as possible reasons. One neglected alternative is that neural degradation of early visual areas underlies greater place-specific dedifferentiation, such as age-related thinning in peripheral regions of primary visual cortex^15^ which is preferentially connected to place-selective regions^16,17^.

To gain mechanistic insights into the neurobiological basis of dedifferentiation, we simulated age-related neural degradation by applying an *in silico* network degradation approach (Fig. 1a) to a neurobiologically inspired model of the ventral visual stream (VVS). We tested whether degradation is sufficient to produce dedifferentiation in model IT, and how degrading different areas produces empirically observed dedifferentiation effects. We then tested whether degrading the model’s peripheral V1 selectively reduces place differentiation in IT, capturing category-specific effects in aging.

**Figure 1:**
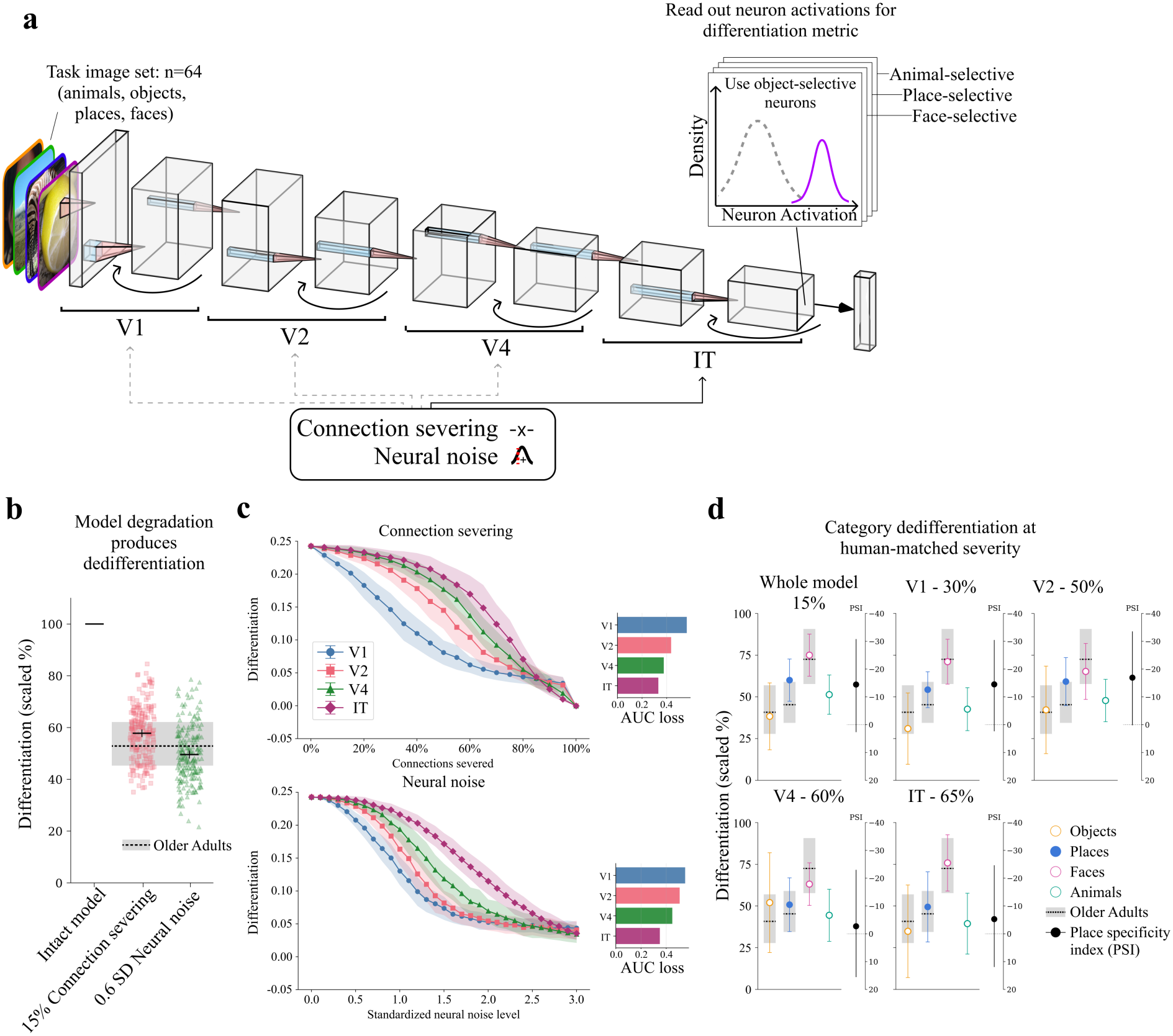
*In silico* network degradation produces aging-like dedifferentiation. **a**, CORnet-RT architecture and degradation methodology. Connections were severed or neural noise was added, either across the whole model or within a single area, and differentiation was read out from category-selective IT neurons. **b**, Overall differentiation after 15% connection severing or neural noise at 0.6 SD (standardized Gaussian)applied across the whole model, scaled to the intact model. Points show individual simulations (*n* = 200); solid black lines show means with error bars indicating bootstrap 95% CIs (10,000 samples). Overall differentiation was averaged over object-, place- and face-selective neurons only, as the human data contain no animal-selective region. The dashed line and gray bands shows the older-adult mean and 95% CIs (scaled to younger adults) based on the fMRI data from Haupt *et al*. [4]. **c**, Overall raw within-between differentiation values in IT across degradation intensities when single model areas were degraded by connection severing (top) or neural noise (bottom). Overall differentiation was averaged over all category-selective neurons (faces, places, animals and objects). Points show means and shading shows *±*1 SD across simulations (*n* = 200). Horizontal bars (right) show area under the curve (AUC) loss for each area. **d**, Category differentiation, scaled to the intact model. For each degraded area, the connection-severing level was set to the level whose overall drop best matched older adults. Markers show means *±*1 SD across simulations. Dashed lines and gray bands show the older-adult means and 95% CIs (scaled to younger adults). Side panels show the place specificity index (PSI) with bootstrap 95% CIs including human data resampling. The PSI axis is inverted, so negative values (places less dedifferentiated than in older adults) appear above zero (Tables S1 and S2).

We used CORnet-RT^18^, a recurrent convolutional neural network (CNN) developed to model the VVS, with hierarchical correspondence^19^ to human V1, V2, V4, and IT (Fig. 1a; see Methods and Fig. S1 for other models). We applied biologically inspired degradation *in silico* by severing or adding neural noise to the connections of the model. Specifically, we degraded connections either across the entire model or selectively within V1, V2, V4, or IT, targeting both inputs to each area and connections between the convolutional layers within it (Fig. 1a). We varied degradation intensity by increasing the percentage of randomly severed connections or the magnitude of added noise. For each intensity and target area, we ran 200 simulations with independently sampled connection damage or noise and averaged differentiation across simulations.

We used stimuli from a human fMRI study of age-related dedifferentiation^4^ and measured differentiation for faces, objects, animals, and places among category-selective neurons in model IT (see Methods), so that model differentiation could be compared directly with empirical findings from category-selective cortex.

Following prior work^4,14^, we defined category differentiation as the mean within-category similarity (Pearson’s *r*) minus the mean between-category similarity of model neuron activations (e.g., mean similarity between each face and other faces minus mean similarity of faces and images in all the other categories). Lower values indicate less distinct representations (ded-ifferentiation).

Our *in silico* network degradation approach captured age-related dedifferentiation. Both connection severing and neural noise reduced differentiation relative to the intact model (Fig. 1b). With 15% connection severing or neural noise at 0.6 SD applied across the entire model, the overall magnitude of dedifferentiation matched that reported by Haupt *et al*. [4] (see Table S1).

When model areas were degraded individually, damage to early areas affected IT differentiation more than damage to later areas (Fig. 1c; see Fig. S1 for other models), suggesting that degradation in early visual areas may contribute to dedifferentiation in IT (see below). We also found dedifferentiation with neuron deletion (Fig. S2), although neuron death in normal aging is unlikely^20^.

To test whether the model reproduced the human pattern of age-related dedifferentiation across categories, we set the degradation intensity so that overall dedifferentiation matched the human data and compared each category with its human counterpart (Fig. 1d; Table S2; see Fig. S3 for a robustness check). Results suggested a trend of places falling short of their human level compared to faces and objects. We summarized this with a place specificity index (PSI): how far places deviate from their human level, minus the mean deviation for faces and objects, where negative values mean that places fall short. Places fell short particularly with whole-model and early-area (V1/V2) degradation (PSI = *−*14.6 to *−*16.8%) and, to a lesser extent, with later-area (V4/IT) degradation (*−*4.5 to *−*5.5%). However, given the high variability in the human data, these differences were not statistically significant (*p ≥* .275, Holm-corrected). This trend is nonetheless notable as places are often the most dedifferentiated category in older adults^3^, raising the possibility that uniform degradation within areas is not sufficient to capture human-like dedifferentiation patterns.

We hypothesize that stronger place dedifferentiation could also arise upstream, from spatially non-uniform degradation to early visual pathways that contribute to place processing. V1 is a natural candidate, as its degradation produced the strongest IT dedifferentiation (Fig. 1c) and it undergoes age-related cortical thinning^21,22^, particularly in its peripheral visual field representation^15^. Notably, peripheral V1 is more functionally and structurally connected to place-selective regions while foveal V1 is more strongly connected to face- and object-selective regions^16,17^. We therefore tested whether peripheral V1 degradation would selectively affect place dedifferentiation by applying degradation to the periphery of model V1, which is organized by visual-field position like human V1, and measured category-specific dedifferentiation in IT.

To test this, we first verified that the model shows the same spatial biases as human visual cortex: peripheral V1 drives place-selective regions, and foveal V1 drives face- and object-selective regions. To do this, we estimated how much V1 neurons contribute to the activation of each category-selective IT neuron by computing a gradient for each V1 neuron and comparing these values to gradients computed using random sets of IT neurons (Fig. 2a). The comparison (Cliff’s delta) at each V1 location can be taken as an estimate of how much peripheral or foveal V1 drive category-selective neurons in IT, with the random control accounting for drive solely from image features.

**Figure 2:**
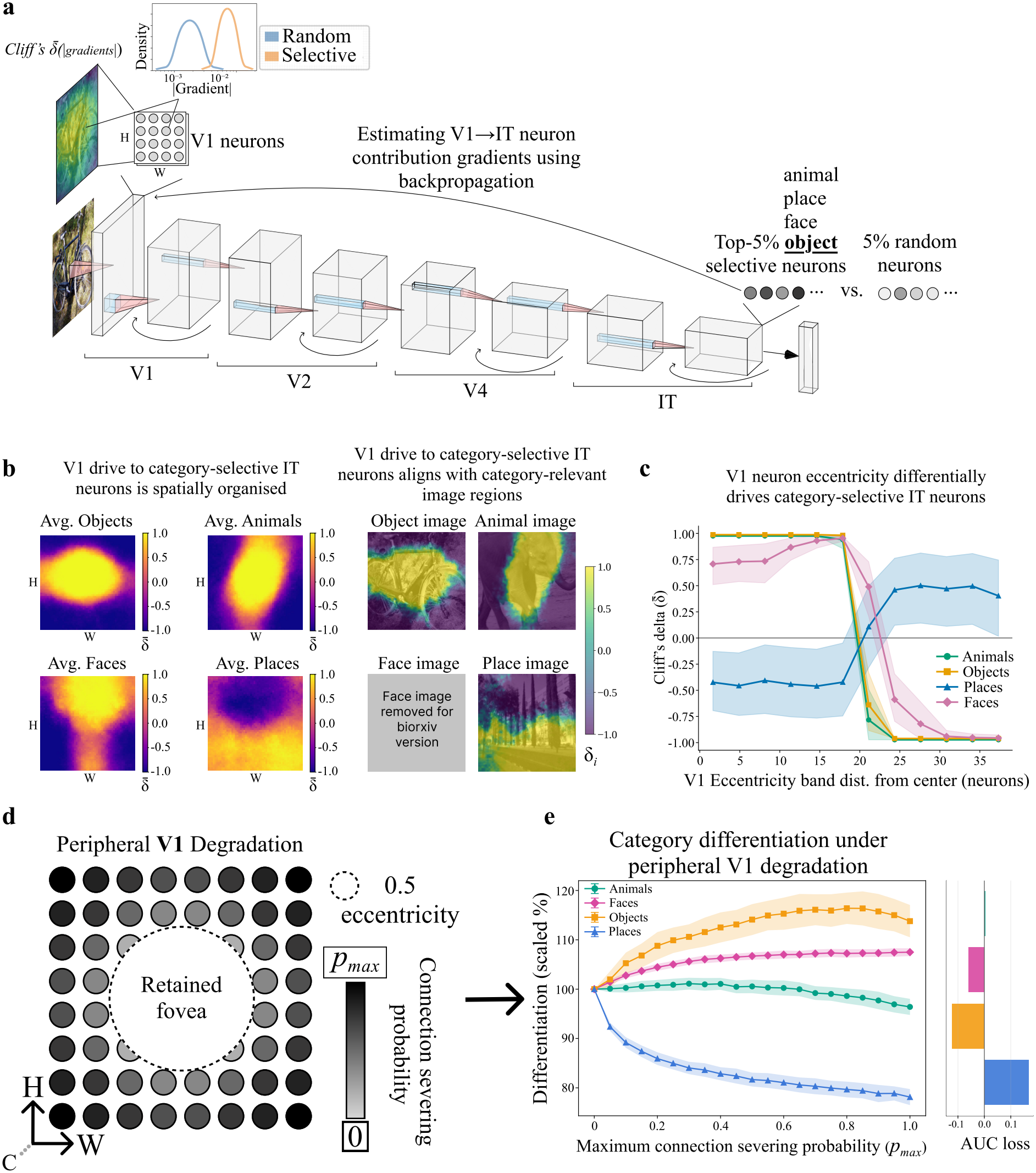
Peripheral V1 degradation captures category-specific dedifferentiation. **a**, Method for assessing retinotopic biases for categories in the model. **b**, Left, Cliff’s delta 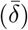 values across V1 neurons for each category for all category images. Right, example images with per image *δ*_*i*_ values overlaid showing which image areas most drive neurons selective for each category. **c**, Cliff’s 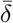 between the gradient density of selective and random IT neurons per category in each eccentricity band. X-axis shows the distance from the V1 center to each eccentricity band’s midpoint. **d**, Peripheral V1 degradation method. **e**, Left, Within-between differentiation (scaled by intact model differentiation) from peripheral V1 damage per category across degradation intensities. Right, AUC loss for each category. In **c** and **e**, points show mean values and **c** shading represents 95% CI, while shading in **e** represents *±*1 SD.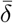: mean Cliff’s delta, *δ*_*i*_: per image Cliff’s delta, H: V1 featuremap height axis, W: V1 featuremap width axis, C: V1 channels, *p*_*max*_: maximum connection severing probability.

As predicted, place-selective IT neurons were driven more by peripheral V1 neurons while face-, object-, and animal-selective IT neurons were driven more by foveal V1 neurons (Fig. 2b). Comparing gradient density for category-selective and random IT neurons across radial eccentricity bands in V1 (see Methods), we found that place-selective neurons showed a peripheral bias, with a greater drive than random IT neurons farther from the center of V1, whereas face-, object-, and animal-selective neurons showed the opposite pattern (Fig. 2c). These biases persisted for images from non-target categories (Fig. S4), indicating that they reflect the neurons’ spatial sensitivity rather than image content.

After confirming the peripheral place-processing bias, we severed the model’s V1 connections with an increasing probability towards the periphery while retaining a central foveal area (Fig. 2d; Methods). This resulted in substantial place dedifferentiation, while differentiation for faces, objects, and animals was relatively preserved across degradation levels (Fig. 2e). Peripheral neuron deletion produced the same results (Fig. S5).

To confirm that this selective place dedifferentiation depends on peripheral–foveal biases, we show that foveal degradation produced the reverse category-level pattern in the model (Fig. S6). These results support our idea that degradation in early visual areas can propagate to category-specific representational changes in higher-level areas, and offer an explanation for the more consistent place dedifferentiation effect in aging. Alongside the strong effect of V1 degradation on dedifferentiation (Fig. 1c), this suggests that degradation in early visual regions plays a crucial role in age-related dedifferentiation.

Together, our analyses show that neural degradation is, in principle, mechanistically sufficient to produce age-related dedifferentiation. Furthermore, degradation in early visual areas produced greater downstream dedifferentiation than equivalent degradation in later areas, likely due to effects compounding across successive processing stages, a property shared by deep networks and the visual hierarchy. Therefore, the degradation underlying dedifferentiation need not occur in the higher-level visual areas where dedifferentiation is observed.

Both neural noise and connection severing induced dedifferentiation in model IT. Consistent with this, neuroimaging studies have associated greater dedifferentiation with both lower gamma-aminobutyric acid (GABA) levels^12^ and reduced white matter tract integrity along the VVS^13^. Reduced GABA levels have, in turn, been linked to increased neural noise in visual cortex^23^ and to dedifferentiation in a computational model^24^. Our results predict that dedifferentiation should depend on GABA levels in *early* visual areas and the integrity of their white matter connections to IT. Because the model lacks top-down feedback, these effects also suggest that dedifferentiation need not reflect age-related changes in top-down attention^25,26^.

Our results provide a mechanistic account of the disproportionate place dedifferentiation observed in aging. Degrading peripheral V1 selectively reduced place differentiation because, as in the human VVS, place-selective IT neurons relied disproportionately on peripheral input. Category-specific dedifferentiation may therefore reflect an interaction between the spatial profile of degradation and how categories are spatially structured in the visual environment.

In sum, dedifferentiation need not stem from uniform or local degradation, but can arise from degradation of specific upstream regions and pathways.

## 2 Methods

### 2.1 Models and Experimental Data

#### 2.1.1 Models

We used CORnet-RT^18^ (5 time-steps), a recurrent CNN, to model the VVS. While earlier models such as AlexNet^27^ and VGG16^28^ already achieve a relatively high level of representational alignment with the human VVS, CORnet-RT achieves this alignment through design choices intended to mimic the brain, including recurrent processing^29^ and a block structure corresponding to brain regions V1, V2, V4, and IT^18^. Each block contains two convolutional layers. The output of the second layer is retained as the block’s internal state and added to the feedforward input of the same layer at the next time-step; because this layer preserves tensor dimensionality, this provides a simple implementation of within-area recurrence. As image-driven activity takes four time-steps to reach IT, five time-steps permit the first recurrent update of an image-driven IT state. In contrast to the architecture of CORnet-RT, many CNN models implement elements that boost performance on their task objective (e.g., object-recognition accuracy) with limited or no gains in alignment with the brain during naturalistic image viewing^30^. Despite its simple and biologically oriented architecture, CORnet-RT nevertheless achieves relatively high ImageNet^31^ object-classification performance, with 55.4% top-1 accuracy and 78.9% top-5 accuracy^32^, and shows alignment with both neural and behavioral responses. This combination of neural and behavioral alignment, architectural simplicity, and solid object-classification performance led us to focus our analyses on CORnet-RT.

To complement the CORnet-RT model and confirm that the results were broadly applicable, we performed the main analyses (Fig. S1) using feedforward models VGG16^28^, AlexNet^27^, and CORnet-RT (4 time-steps; feedforward setting), as well as a recurrent CORnet variant, CORnet-S^18^. For AlexNet and VGG16, we partitioned the model areas using their five-block structure, so that degradation was applied to a single block at a time. AlexNet contains five convolutional layers, one within each block, whereas VGG16 contains 13 convolutional layers, with two or three layers per block. Both models contain three fully connected layers after the convolutional blocks. Throughout the analyses, a connection refers to an individual learned convolutional weight, whereas a filter refers to the complete set of weights and the bias parameter producing one output feature channel. All models were pre-trained on ImageNet’s 1,000-way object-classification task^31^. The AlexNet and VGG16 models were loaded from Torchvision^33^, while the CORnet models were implemented using the latest version of the official CORnet GitHub repository (https://github.com/dicarlolab/CORnet; downloaded 10 February 2025).

#### 2.1.2 Experimental Stimuli

To allow direct comparison with human data, we used the image set from a previous functional magnetic resonance imaging (fMRI) study of age-related dedifferentiation^4^. The image set comprises 16 images from four categories (64 total): animals, objects, faces, and places (image-set download: https://osf.io/xeukw). Animals, objects, and faces were clearly depicted, centrally positioned, and occupied most of the image frame. The animals included mammals and birds, while the faces varied in age, sex, and ethnicity. The objects included fruit, vegetables, other food items, transport, and other everyday objects. Places depicted a range of outdoor natural and built environments, including mountains, trees, houses, and other buildings. The images were originally obtained from COCO^34^, FairFace^35^, and Places-365^36^.

#### 2.1.3 Human Reference Data

We compared model dedifferentiation with the fMRI results of Haupt et al.^4^. They reported within-between differentiation for 21 younger and 21 older adults in the fusiform face area

(FFA), lateral occipital complex (LOC), and parahippocampal place area (PPA), each computed for the region’s preferred category against each of the other three categories (e.g., PPA place vs. face). We used their results in normalized space with responses averaged over all runs (their Fig. 6B and Table S5). To keep the participant-level data of Haupt et al. unseen for a planned registered report, we did not access the openly available dataset and instead digitized the individual participant values plotted in their Fig. 6B. We shifted the digitized values so that their mean equaled the mean reported in their Table S5, which retained the plotted spread while making the human point estimates identical to the published means. Because the human data contain no animal-selective region, animals were excluded from all model-human comparisons.

### 2.2 Identification of Category-Selective Neurons

To quantify dedifferentiation, we identified category-selective neurons using an approach analogous to the functional-localizer procedures employed in human fMRI studies^4^. Whereas these studies define category-selective regions of interest by selecting voxels that respond most strongly to a target visual category, we defined corresponding model populations by selecting the neurons with the strongest responses to that category within the most category-selective model layer.

To measure category selectivity while limiting reliance on low-level, stimulus-specific visual features, we first collected an independent localizer image-set. For each of the four categories, 100 images were initially collected from the royalty-free stock-photo database, Un-splash (https://unsplash.com/) using the following search terms: face, “face”; place, “scene” and “house”; object, “tool”, “clothing”, “fruit”, “vehicle”, and “object”; animal, “animal” and “mammal”, matching many of the subcategories found in the experimental stimuli. From each initial set, we selected the 20 most representative exemplars based on minimal clutter, naturalistic color grading, and clear membership to the target category.

We passed these 80 independent images (20 per category) through each intact CNN model and compared category selectivity across candidate convolutional layers and fully connected layers, using activations after the activation function. Post-activation values were used to in-corporate the non-linear, non-negative aspects of neuronal processing. For every neuron and category, we used a Mann–Whitney *U* -test^37^ to compare the neuron’s responses to the 20 target-category images with its responses to the 60 images from the remaining categories. We used this non-parametric test because it does not assume normally distributed activations and is robust against unequal sample sizes. The test measures whether values from one distribution tend to be larger or smaller than values from the other distribution. For each candidate layer, we averaged *U* among the top 5% of neurons within each category and then across the four categories. This criterion selected the IT block output for the CORnet models, the post-ReLU output of the first fully connected layer for AlexNet (classifier.2), and the post-ReLU output of the second fully connected layer for VGG16 (classifier.4). The subsequent selectivity and differentiation analyses were conducted within these most category-selective layers.

Within the selected layer of each model, the same Mann–Whitney procedure was used to identify the category-selective neurons. The four *U* -values obtained for each neuron described its selectivity for faces, places, animals, and objects, respectively. For each category, neurons were ranked by their *U* -values and the top 5% within the selected layer were retained as that category’s selective population.

The same four groups of top 5% category-selective neurons were used throughout all sub-sequent analyses. They were defined once using the intact model and independent localizer images, and were then held constant across all degradation conditions. Thus, for example, face differentiation was always computed from the same face-selective neurons rather than from neurons reselected after each degradation. The 5% threshold was chosen to balance high category selectivity within the localizer images with generalization of that selectivity to the independent experimental images. For example, in analyses not reported here, a top 1% threshold in AlexNet produced lower within-between differentiation for the experimental images than the top 5%, suggesting that the narrower population did not generalize as well beyond the localizer set. Analyses using the top 1% and 10% nevertheless produced qualitatively similar results in the main analyses (Fig. S7).

### 2.3 Quantifying Representational Differentiation

We quantified category differentiation using the within-between differentiation metric, a measure of representational category distinctiveness commonly used in human fMRI studies^14^. For each randomized degradation simulation (i.e., a specific degradation type, target area, and intensity), we extracted activation vectors for all 64 experimental images from the selected model layer. For a target category *c*, each vector included only the top 5% of neurons selective for that category.

We then constructed a 64 *×* 64 representational similarity matrix (RSM) by calculating Pearson’s *r* between these activation vectors for every pair of images.

Category differentiation was the mean correlation between pairs of images from the target category minus the mean correlation between target-category images and images from the other three categories:

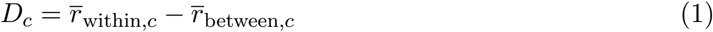

For example, we calculated face differentiation from the RSM based on face-selective neurons by subtracting the mean face-place, face-object, and face-animal correlations from the mean face-face correlation. Higher values therefore indicate that images from the target category have more similar activation patterns to one another than to images from other categories, whereas lower values indicate reduced representational distinctiveness. We computed this measure separately for faces, places, animals, and objects. Their mean,

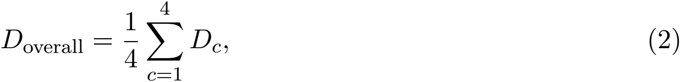

provided the overall differentiation measure used to compare degradation types and model areas.

#### 2.3.1 Scaled Differentiation and Area Under the Curve

To compare categories and degradation conditions on a common scale, we expressed differentiation relative to the intact model. Scaled differentiation of category *c* at degradation intensity *x* was

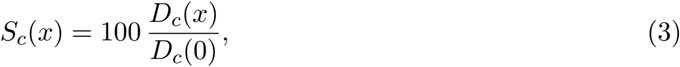

where *D*_*c*_(0) is the differentiation of the undamaged model, so that 100% indicates no dediffer-entiation. In comparisons with human data, which contain no animal-selective region, overall scaled differentiation was the mean of *S*_*c*_ across faces, objects, and places.

To summarize the effect of a degradation condition across all tested intensities, we calculated a normalized area under the curve (AUC) loss. At intensity *x*, the loss of differentiation relative to the intact model was

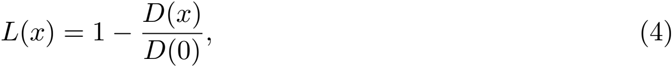

where *D*(*x*) is the mean differentiation across the 200 simulations at intensity *x*, either overall (Eq. 2) or for a single category (Eq. 1), and *D*(0) is the corresponding differentiation in the undamaged model. We integrated *L*(*x*) across the tested degradation range using the trapezoidal rule and divided the result by the width of that range:

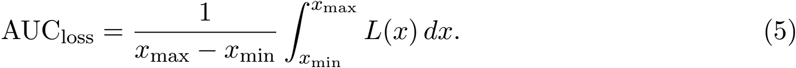

### 2.4 Simulating Neural Degradation

To investigate which biological processes might underlie age-related dedifferentiation, we implemented four distinct types of artificial degradation to the models. Three of these were adapted from prior work in connectionist neuropsychology (e.g.,^38,39^) to simulate connection severing, neuron deletion, and neural noise. The fourth was a novel implementation of peripheral V1 degradation inspired by biological aging patterns in the primary visual cortex^15^.

#### 2.4.1 General Degradation Procedure

For each of the three spatially uniform manipulations, we applied degradation to the entire model or one model area at a time (V1, V2, V4, or IT for CORnet-RT). We varied the intensity from an intact model to the maximum value specified below for that manipulation. At each combination of degradation type, target area, and intensity, we ran 200 independent simulations. Each simulation used a fresh random draw, so that a new subset of connections or neurons was selected for deletion, or a new sample of noise was generated. The 64 experimental stimuli were passed through each degraded model, and activations from the fixed category-selective populations were saved to calculate differentiation. The V1 peripheral degradation was applied only to V1 and otherwise followed the same 200-simulation procedure.

#### 2.4.2 Connection Severing and Neuron Deletion

To simulate the disruption of white-matter connectivity in the aging VVS, we used the connection-severing method from classical lesioning studies^40^. For each simulation, a proportion of random convolutional weights in the target area was set to zero separately within every convolutional layer. Degradation ranged from 0% to 100% of weights in 5-percentage-point increments. Because the first convolutional layer of an area receives the output of the preceding area, this manipulation affected both incoming and within-area connections. For example, when V4 was targeted, connection severing removed a random subset of the connection weights carrying signals from V2 to V4 as well as connection weights between convolutional layers within V4.

Although neuron deletion has a less direct biological analogue in healthy aging, where neuronal cell death is relatively rare^20^, we included it to make our analyses comparable with prior lesion-based modeling studies^38–40^. For each simulation, a proportion of random convolutional filters were selected separately within every convolutional layer of the target area. We set all learned weights and the bias parameter of each selected filter to zero, causing the direct output of that convolutional filter to be zero. The deleted proportion ranged from 0% to 100% of filters in 5-percentage-point increments. We also tried deleting the output activations of randomly selected filters instead of effectively deleting entire feature channels and obtained qualitatively similar results.

#### 2.1.3 Neural Noise

To simulate the effects of age-related increases in neural noise, we added zero-mean Gaussian noise to the convolutional weights within a target area. Because the magnitude of learned weights varies across layers, a fixed noise distribution would not represent an equivalent perturbation throughout the model: the same absolute noise would be proportionally larger in layers with smaller weights. We therefore matched the scale of the noise to the weight distribution of each layer. For layer *l*, each noise value was sampled independently as

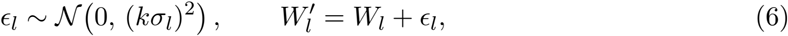

where *N* denotes a Gaussian (normal) distribution (*mean*=0, *SD* =*kσ*_*l*_), *σ*_*l*_ is the standard deviation of the intact weights in layer *l, k* controls noise intensity, and 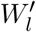denotes the resulting perturbed weight values for that layer. This scaling made the noise comparable in relative magnitude across layers and model areas. The scaling factor *k* ranged from 0 to 3 in increments of 0.1.

### 2.5 Comparison with Human Dedifferentiation

#### 2.5.1 Relative Drop in Differentiation

To compare model and human dedifferentiation on a common scale, we expressed both as a relative drop in differentiation (Table S1). For the model, the relative drop of category *c* was ∆_*c*_(*x*) = 100 *− S*_*c*_(*x*) (Eq. 3), computed within each simulation, so that 0% indicates no dedif-ferentiation. For humans, the relative drop of each region was computed from the group means of its three comparisons,

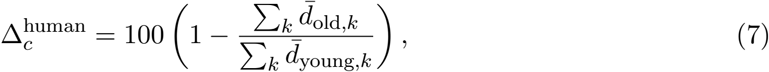

where 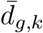 is the mean differentiation of age group *g* for comparison *k* (e.g., for PPA, place vs. face, place vs. object, and place vs. animal). We used this ratio of summed group means because younger and older adults are different participants, so no per-participant drop exists. Overall dedifferentiation was the mean of the face, object, and place drops, computed within each simulation for the model.

#### 2.5.2 Place Specificity Index

At the tested degradation level whose overall drop was closest to the human overall drop (Table S1), we computed each category’s deviation from its human level, 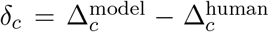 (percentage points). We then defined the place specificity index (PSI) as

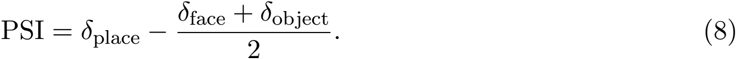

A PSI of zero indicates that places were as close to their human level as faces and objects were to theirs, whereas negative values indicate that places were less dedifferentiated than in older adults relative to the other categories. Confidence intervals and two-sided p values were obtained from a percentile bootstrap (100,000 resamples) that resampled both the model simulations and the human participants, and p values were Holm-corrected across the five degraded areas.

#### 2.5.3 Place Slope Difference

As a robustness check that does not rely on the single matched intensity from the PSI (Fig. S3; Table S2), we examined how quickly each category approached its human level as degradation increased. For each category and tested intensity, progress was defined as the mean model drop divided by the human drop of that category (1 = human level). Progress was regressed on the mean overall drop by ordinary least squares across the tested intensities up to and including the first that reached the human overall drop. This range restricted the fit to the severities relevant for the human comparison. Slopes were expressed per 10 percentage points of overall drop. The place slope difference was the place slope minus the mean of the face and object slopes; negative values indicate that places approached their human level more slowly than faces and objects as degradation increased. Confidence intervals and Holm-corrected p values were obtained with the same bootstrap as for the PSI, refitting the slopes in each resample.

### 2.6 Peripheral V1 Analyses

We performed two related analyses to test whether stronger place dedifferentiation in aging could arise from peripheral V1 degradation. Testing this in a model is meaningful only if it first reproduces the organization of the human VVS: peripheral V1 drives place-selective regions, and foveal V1 drives face- and object-selective regions. We therefore first assessed whether category-selective IT neurons in CORnet-RT depended preferentially on foveal or peripheral parts of V1 (Fig. 2a-c), and then degraded the model’s peripheral V1 and measured the resulting category-specific differentiation in IT (Fig. 2d-e).

#### 2.6.1 Gradient-Based V1-to-IT Sensitivity Analysis

To test whether CORnet-RT reproduced the category-specific IT-V1 biases observed in the human VVS, we used a gradient-based sensitivity analysis to quantify how strongly category-selective IT neurons depended on activations across the V1 visual field. We performed this analysis on the activations after the input convolution and its nonlinear activation function in V1 (V1.nonlin input), because this layer provides the most direct neuron-to-pixel mapping within the V1 area of the model and includes the activation function of the first convolutional layer. For a target category (e.g., faces), we used backpropagation to measure how strongly each V1 layer neuron influenced each category-selective neuron in IT, using the absolute gradient magnitude to capture sensitivity regardless of whether the influence increased or decreased the IT response. For each of the 16 target-category images, this procedure was repeated for every neuron in the top 5% category-selective IT population. We then averaged the absolute gradients across selective IT neurons, yielding one mean V1 gradient tensor per image, organized by feature channel and spatial position.

We compared these category-selective gradients with gradients from randomly selected IT neurons to determine whether an apparent spatial bias exceeded that expected from the image set and the model’s general architecture. The entire procedure was repeated 100 times using random sets containing the same number of IT neurons as the category-selective population. Within each random set, gradients were again averaged across IT neurons, leaving 100 equally sized control populations for every image.

Because we were specifically interested in which parts of V1 were driving each IT population, we averaged each gradient tensor across the 64 V1 feature channels and divided the resulting spatial map by its sum. Each value therefore reflected the share of the population’s total V1 sensitivity at that position. At each spatial position *x*, we compared the selective map with each random-control map of the same image and averaged the result across images:

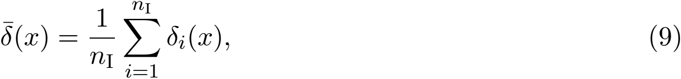

where *δ*_*i*_(*x*) is Cliff’s delta between the normalized selective value at position *x* of image *i* and those of the 100 random-control sets on the same image (i.e., the proportion of random sets with a lower value minus the proportion with a higher value), and *n*_I_ = 16 (number of images). Both *δ*_*i*_(*x*) and 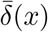 range from -1 to 1. Positive values indicate that the category-selective population allocated a larger share of its V1 sensitivity to that position than equally sized random populations, values near zero indicate no difference, and a value of 1 indicates that the selective population exceeded every random set on every image. Because images and random sets were the sampled units, and positions were not treated as samples, we estimated pointwise 95% confidence intervals by independently resampling images and random sets with replacement over 10,000 bootstrap repetitions. Repeating this analysis for all four categories produced the category-specific spatial bias maps shown in Fig. 2b. We also repeated each category-selective analysis separately with the 16 images of each of the three non-target categories. These maps showed similar spatial patterns, indicating that these biases were not driven by low-level features unique to the target-category images (Fig. S4).

#### 2.6.2 Retinotopic Bias by Eccentricity

To quantify the computed category-specific spatial biases along the foveal-peripheral axis (Fig. 2c), we reduced each normalized two-dimensional spatial gradient map described above to a one-dimensional radial profile. Because the maps were normalized, these profiles describe how each population’s V1 sensitivity was distributed from the fovea to the periphery, independent of its overall magnitude.

We defined the eccentricity of each point in the V1-derived spatial gradient map as its Euclidean distance from the center. The range from the center to the corners was divided into 12 equal-width radial bins. For each image, we summed the normalized gradients within each bin and divided this value by the number of values in that bin, producing an area-normalized gradient-density value for each eccentricity band. We applied the same procedure separately to each of the 100 random-control sets. This yielded 16 selective image profiles and 1,600 random-control profiles (100 sets *×* 16 images) for each category.

For each eccentricity band *b*, we computed 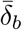 as for the spatial maps (Eq. 9), comparing each image’s selective band density with those of the 100 random-control sets for the same image and averaging across the 16 images. Positive values indicate a larger share of normalized gradient density for category-selective than for random IT neurons at that eccentricity, whereas values near zero indicate no difference. The shaded regions in Fig. 2c show pointwise 95% bootstrap confidence intervals, obtained as for the spatial maps.

#### 2.6.3 Peripheral V1 Degradation

We next tested whether the greater place dedifferentiation reported in healthy aging could arise from spatially biased, peripheral degradation in V1. Spatial locations near the center of the image were preserved, while locations farther towards the periphery had an increasingly high probability of deletion (Fig. 2d). The manipulation was applied to the input convolution in model V1 (V1.conv input) in CORnet-RT.

To implement this pattern, we assigned every spatial output location of the V1 convolution an eccentricity value based on its Euclidean distance from the center, normalized from 0 at the center to 1 at the corners. Standard convolution applies the same learned filter weights for every output location, a process known as weight sharing. We therefore computed the convolution independently for each spatial location for each filter, allowing a given weight to be retained when its filter was applied at one location but deleted when the same filter was applied at another. At each location, every connection weight within each output filter was independently retained or set to zero according to the location’s eccentricity-dependent deletion probability. Motivated by the pattern of age-related cortical thinning in human V1^15^, weights contributing to locations with eccentricity at or below 0.5 were retained. Beyond this boundary, deletion probability increased linearly with eccentricity, from zero at 0.5 to a maximum probability *p*_max_ at the corners:

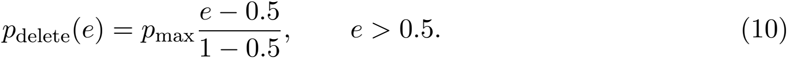

For each degradation simulation, the location-specific weight-deletion pattern was sampled once and held fixed across all images and recurrent time steps. The intensity of peripheral degradation was varied by increasing *p*_max_ from 0 to 1 in increments of 0.05. Retaining central regions defined by eccentricity thresholds of 0.1 or 0.3 produced qualitatively similar stronger place ded-ifferentiation (Fig. S8). We additionally applied the same eccentricity-dependent deletion profile directly to V1 unit activations, independently setting activations at selected spatial locations across feature channels to zero, and again obtained qualitatively similar category-specific results (Fig. S5).

## 3 Data availability

All data to reproduce the main and supplementary figures are available on Zenodo at https://doi.org/10.5281/zenodo.22864336.

## 4 Code availability

Scripts for running all the simulations and analysis presented in the paper are available via GitHub at https://github.com/AkillesU/dedifferentiation-models

## 5 Acknowledgements

This work was supported by the UKRI Economic and Social Research Council through the South East Doctoral Training Arc (SEDarc) [grant ES/Y001656/1 to A.R.].

