## Supplementary materials for "A Mechanistic Account of Dedifferentiation in the Aging Visual System"

### 6 Supplementary Materials

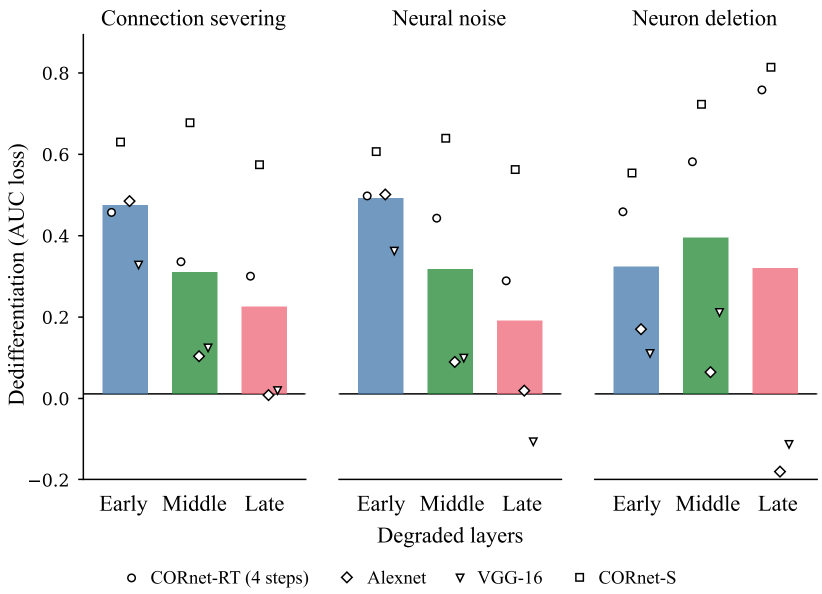


**Figure S1: *In silico* network degradation shows early-to-late dedifferentiation gradient with connection severing and neural noise.**
To complement the analyses shown in the main text, we ran the same experiment using other CNN models: AlexNet, VGG-16, CORnet-RT (4 time-steps), CORnet-S. For the CORnet models, degradation was applied to areas V1, V2, V4, and IT while for AlexNet and VGG-16, degradation was applied to blocks 1-5. We computed differentiation from the top-5% category-selective neurons for all models (see Methods). The plot shows dedifferentiation from neural noise, connection severing, or neuron deletion as AUC loss, averaged first across degradation areas within each model and then across models, when degradation is applied to early (block1/V1), middle (block2-3/V2 and V4), or late (block4-5/IT) layers. Points show mean values for individual models.

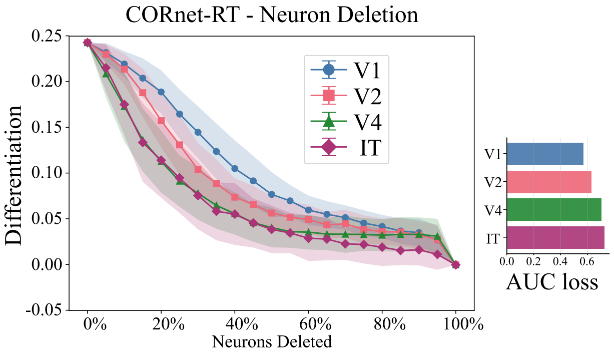


**Figure S2: Differentiation from neuron deletion in CORnet-RT.**
We applied neuron deletion to the model by setting all the parameters of individual convolutional filters in a target area to 0. In practice this sets all the output activations of a given filter to 0 and inhibits any information from passing on. The plot shows raw differentiation in IT shown across degradation areas and intensity. Points show mean values and shading represents $\pm 1$ SD. Horizontal bar plots on the right show area under curve (AUC) loss for each area across degradation intensities.

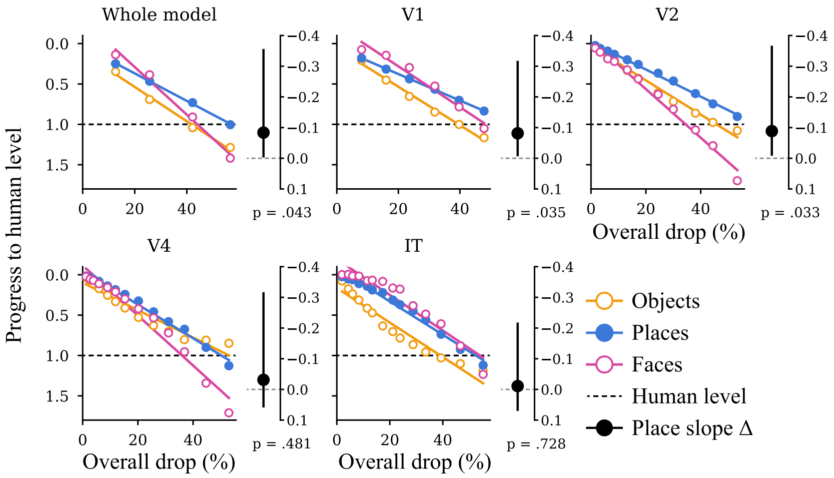


**Figure S3: Places show a trend of approaching their human level of dedifferentiation more slowly under uniform degradation, related to Fig. 1d.**
Progress towards the human level (model drop divided by human drop; dashed line at 1 = human level) for faces, objects, and places, plotted against the model's overall drop for each degraded area. The y-axis is inverted so that greater dedifferentiation moves downwards. Points show means across simulations (n = 200) at each tested degradation level up to the first that reached the human overall drop (excluding the intact model). Lines show ordinary least squares fits. Side panels show the place slope difference (place slope minus the mean face and object slope, per 10 percentage points of overall drop) with 95% bootstrap CIs (100,000 resamples) and uncorrected two-sided p values (see Table S2 for corrected).

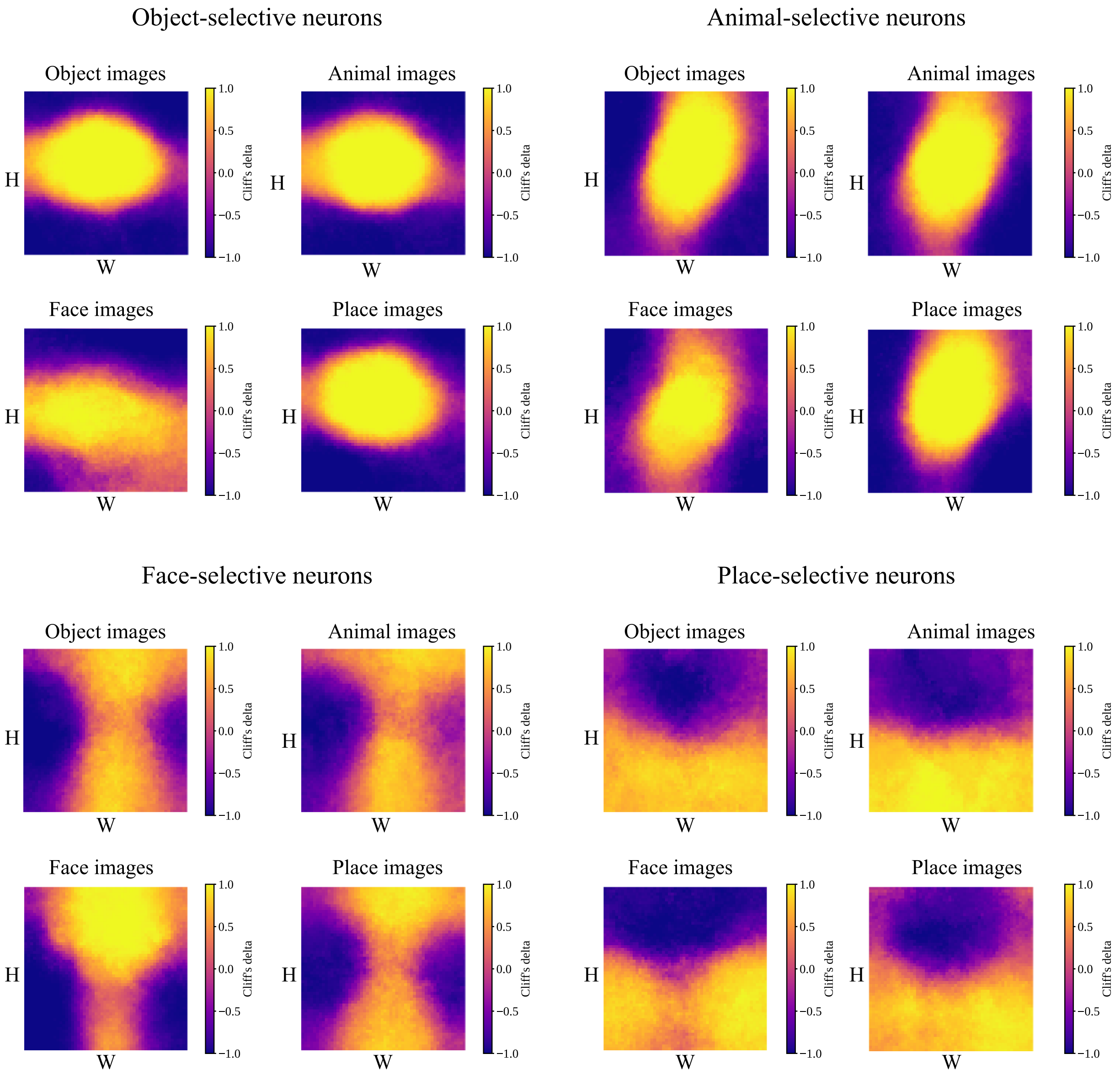


**Figure S4: Category-selective retinotopic biases do not depend on low-level image properties, related to Fig. 2b.**
To test whether the retinotopic biases of category-selective IT neurons were driven by low-level properties of their target-category images, we repeated the analysis for each neuron group using images from the three non-target categories (see Methods). For each group of category-selective IT neurons, a separate plot is shown for each image category. The plots use the same Cliff's delta ($\bar{\delta}$) metric as was used in Fig. 2b. H: V1 featuremap height axis, W: V1 featuremap width axis.

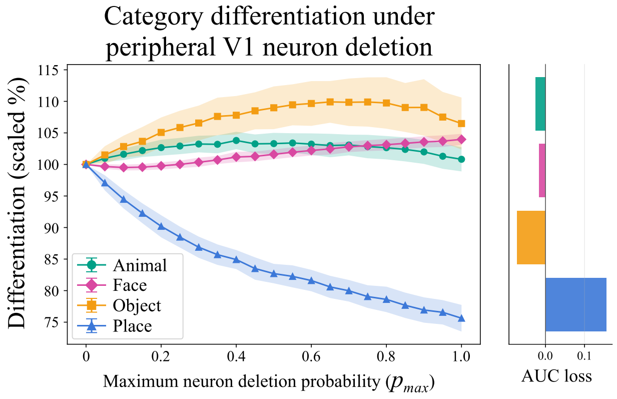


**Figure S5: Peripheral V1 neuron deletion shows similar results to peripheral connection severing.**
We also ran the peripheral V1 degradation analysis by deleting entire neurons instead of connections in the same V1 layer. Neurons were deleted by eccentricity with a retained fovea up to 0.5 standardized eccentricity, akin to the connection severing version. The plot shows scaled differentiation per category percentage-scaled to the undamaged model baseline with points representing mean values for a category at each degradation intensity. The shading shows ±1 SD. Horizontal bar plots on the right show area under curve (AUC) loss for each category across degradation intensities.

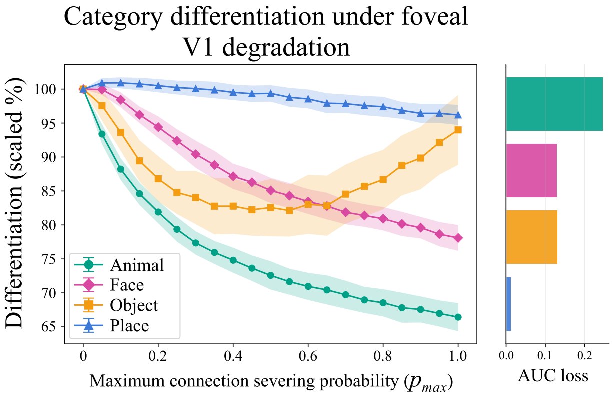


**Figure S6: Foveal V1 degradation retains place differentiation, related to Fig. 2e.** To support the claim that peripheral biases in V1 were driving category-specific dedifferentiation we degraded the V1 fovea from 0.5 eccentricity to the center with increasing probability. The method used was a reverse version of the peripheral V1 degradation with the peripheral neurons kept intact. The plot shows scaled differentiation per category percentage-scaled to the undamaged model baseline with points representing mean values for a category at each degradation intensity. The shading shows ±1 SD. Horizontal bar plots on the right show area under curve (AUC) loss for each category across degradation intensities.

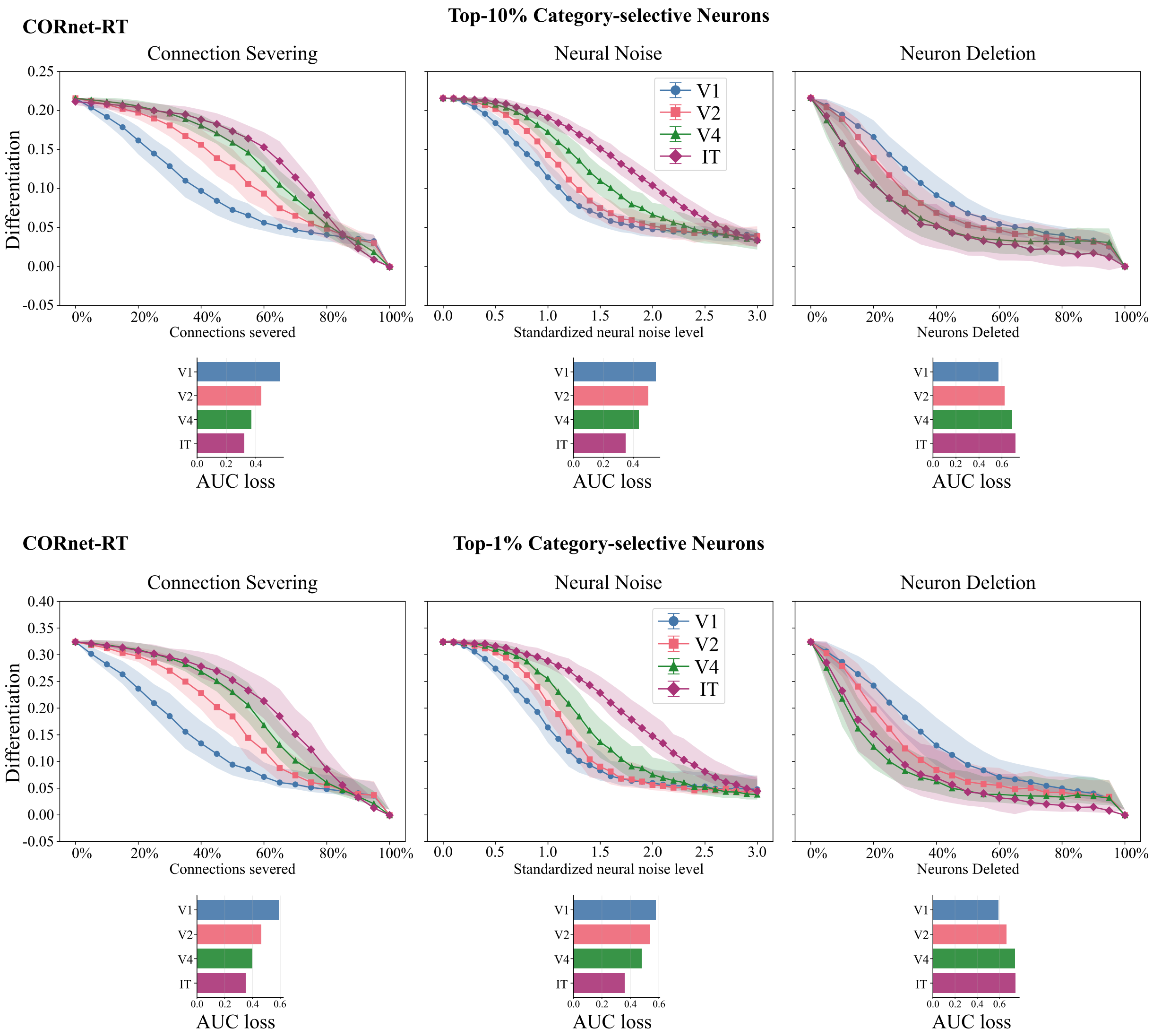


**Figure S7: Changing category-selectivity thresholds for IT neurons in CORnet-RT does not qualitatively change results.**

We applied the same degradation to CORnet-RT as in the main text and used 10 % (top) or 1% (bottom) of the most category-selective neurons in IT to compute differentiation. Changing the fraction of selective neurons used shifts the absolute differentiation values but does not qualitatively affect the pattern of results. The plots show differentiation with points showing mean values and shading representing $\pm 1$ SD. Horizontal bar plots on the right show area under curve (AUC) loss for each region across degradation intensities.

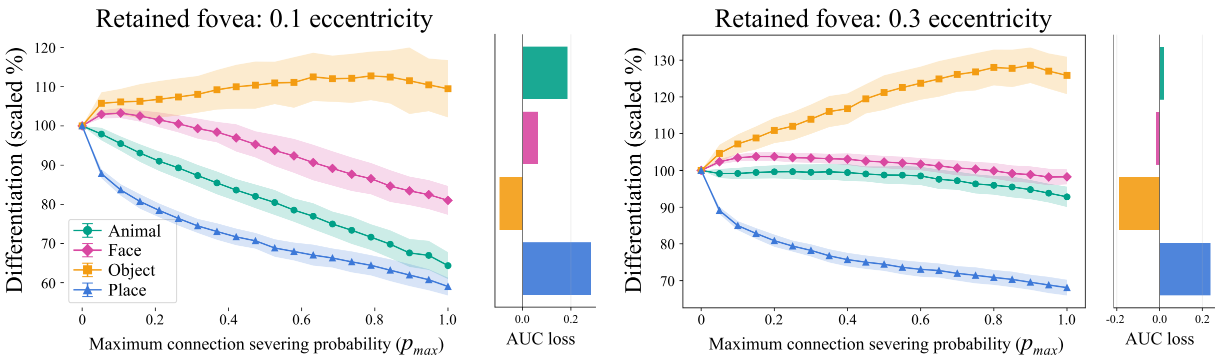


**Figure S8: Other retained fovea values show similar stronger place dedifferentiation, related to Fig. 2e.**
The plots show peripheral V1 degradation with connections in V1 within 0.1 (left) and 0.3 (right) standardised eccentricity from the center of V1 being retained. Differentiation is percentage-scaled to the undamaged model baseline for each category with points representing the mean values for categories at each degradation intensity. The shading shows ±1 SD. Horizontal bar plots on the right show area under curve (AUC) loss for each category across degradation intensities.

**Table S1: Relative dedifferentiation in older adults and in the model.**

| | | | Relative drop in differentiation, % [95% CI] | | | |
|---|---|---|---|---|---|---|
| Source | Area | Level | Overall | Faces | Objects | Places |
| Older adults[4] | | | 47.1 [37.9, 54.7] | 27.5 [9.3, 42.3] | 59.2 [43.0, 72.2] | 54.7 [41.0, 65.6] |
| Connection severing | Whole model | **15%** | **42.2 [40.8, 43.6]** | **25.0 [23.3, 26.7]** | **61.6 [58.9, 64.4]** | **40.0 [38.2, 41.7]** |
| | | 20% | 56.7 [55.3, 58.1] | 39.0 [37.1, 40.9] | 76.1 [73.1, 79.1] | 55.0 [53.2, 56.7] |
| | V1 | 25% | 39.7 [38.4, 41.1] | 21.5 [20.0, 23.1] | 59.3 [56.7, 61.9] | 38.4 [36.7, 40.1] |
| | | **30%** | **47.9 [46.3, 49.4]** | **28.9 [27.0, 30.8]** | **69.0 [66.0, 71.9]** | **45.7 [44.2, 47.2]** |
| | V2 | **50%** | **44.5 [42.7, 46.2]** | **34.7 [32.4, 37.1]** | **57.8 [54.2, 61.4]** | **40.8 [38.8, 42.8]** |
| | | 55% | 53.3 [51.5, 55.1] | 46.7 [44.3, 49.0] | 63.8 [60.1, 67.5] | 49.4 [47.2, 51.6] |
| | V4 | **60%** | **44.7 [42.9, 46.5]** | **36.9 [35.1, 38.6]** | **48.0 [43.8, 52.1]** | **49.2 [47.0, 51.5]** |
| | | 65% | 53.0 [51.0, 54.9] | 47.0 [45.3, 48.6] | 50.2 [45.2, 55.2] | 61.6 [59.5, 63.7] |
| | IT | **65%** | **46.6 [44.5, 48.8]** | **24.2 [21.9, 26.5]** | **65.2 [61.3, 69.0]** | **50.5 [47.7, 53.5]** |
| | | 70% | 55.2 [53.4, 57.1] | 33.9 [31.7, 36.2] | 70.4 [66.8, 74.1] | 61.3 [58.8, 63.8] |
| Neural noise | Whole model | 0.5 SD | 37.8 [36.5, 39.0] | 20.7 [19.2, 22.3] | 55.7 [53.1, 58.3] | 36.8 [35.0, 38.6] |
| | | **0.6 SD** | **50.4 [48.9, 51.9]** | **33.9 [32.0, 35.8]** | **66.6 [63.6, 69.6]** | **50.8 [49.0, 52.5]** |
| | V1 | 0.8 SD | 40.9 [39.6, 42.2] | 21.0 [19.6, 22.4] | 62.6 [60.2, 65.1] | 39.0 [37.4, 40.7] |
| | | **0.9 SD** | **48.1 [46.7, 49.4]** | **26.3 [24.7, 27.9]** | **72.5 [70.1, 74.9]** | **45.4 [43.8, 47.1]** |
| | V2 | 1.1 SD | 41.4 [39.7, 43.1] | 34.2 [32.0, 36.4] | 54.3 [51.1, 57.6] | 35.6 [33.5, 37.7] |
| | | **1.2 SD** | **49.1 [47.5, 50.8]** | **44.3 [42.1, 46.4]** | **56.9 [53.3, 60.4]** | **46.3 [44.2, 48.3]** |
| | V4 | 1.3 SD | 42.1 [40.3, 43.9] | 31.1 [29.2, 32.9] | 50.8 [46.4, 55.0] | 44.5 [42.4, 46.7] |
| | | **1.4 SD** | **48.7 [46.8, 50.6]** | **39.0 [37.2, 40.7]** | **54.3 [49.4, 59.2]** | **52.9 [50.7, 55.1]** |
| | IT | 1.7 SD | 44.2 [42.8, 45.6] | 23.8 [22.0, 25.5] | 56.0 [52.7, 59.2] | 52.8 [51.0, 54.5] |
| | | **1.8 SD** | **48.5 [47.2, 49.8]** | **27.7 [26.0, 29.4]** | **59.5 [56.5, 62.5]** | **58.4 [56.6, 60.1]** |

Relative drop in category differentiation (0% = no dedifferentiation) for older adults (Haupt et al.; FFA for faces, LOC for objects, PPA for places) and for the model at the two tested degradation levels above and below the human overall drop. Overall is the mean of the face, object, and place drops. Bold rows mark the level numerically closest to the human overall drop (shown in Fig. 1d for connection severing). Model values are means across 200 simulations. 95% CIs are percentile bootstrap intervals from 100,000 resamples.

**Table S2: Place specificity index and place slope difference under uniform degradation.**

| | | | Place specificity index (pp) | | | Place slope Δ | | |
|---|---|---|---|---|---|---|---|---|
| Degradation | Area | Level | Estimate [95% CI] | $p$ | $p_{\text{Holm}}$ | Estimate [95% CI] | $p$ | $p_{\text{Holm}}$ |
| Connection severing | Whole model | 15% | −14.7 [−31.1, 2.4] | .089 | .357 | −0.084 [−0.356, −0.003] | .043 | .163 |
| | V1 | 30% | −14.6 [−31.0, 2.5] | .091 | .357 | −0.081 [−0.318, −0.007] | .035 | .163 |
| | V2 | 50% | −16.8 [−33.3, 0.4] | .055 | .275 | −0.088 [−0.367, −0.008] | .033 | .163 |
| | V4 | 60% | −4.5 [−21.2, 12.6] | .583 | 1.000 | −0.031 [−0.317, 0.059] | .481 | .963 |
| | IT | 65% | −5.5 [−22.0, 11.7] | .513 | 1.000 | −0.011 [−0.218, 0.070] | .728 | .963 |
| Neural noise | Whole model | 0.6 SD | −10.9 [−27.3, 6.2] | .203 | .608 | −0.054 [−0.282, 0.023] | .155 | .465 |
| | V1 | 0.9 SD | −15.3 [−31.7, 1.8] | .076 | .359 | −0.057 [−0.245, 0.014] | .105 | .420 |
| | V2 | 1.2 SD | −15.6 [−32.2, 1.5] | .072 | .359 | −0.095 [−0.383, −0.014] | .023 | .116 |
| | V4 | 1.4 SD | −5.1 [−21.7, 12.1] | .545 | 1.000 | −0.047 [−0.301, 0.034] | .238 | .476 |
| | IT | 1.8 SD | 3.4 [−13.0, 20.5] | .689 | 1.000 | 0.018 [−0.167, 0.101] | .744 | .744 |

The place specificity index (PSI) is the place deviation from its human level minus the mean face and object deviation, at the degradation level most closely matching the human overall drop (see Table S1). Negative values indicate that places were less dedifferentiated than in older adults. The place slope Δ is the difference in how quickly places, compared with faces and objects, approached their human level as degradation increased (per 10 percentage points of overall drop; Fig. S3). 95% CIs are percentile bootstrap intervals from 100,000 resamples. $p$ values are two-sided; $p_{\text{Holm}}$ is Holm-corrected across the five degraded areas.
